# Enterovirus-cardiomyocyte interactions: impact of terminally deleted genomic RNAs on viral and host functions

**DOI:** 10.1101/2022.09.15.508200

**Authors:** Alexis Bouin, Michelle N. Vu, Ali Al-Hakeem, Genevieve P. Tran, Joseph H.C. Nguyen, Bert L. Semler

## Abstract

Group B enteroviruses, including coxsackievirus B3 (CVB3), can persistently infect cardiac tissue and cause dilated cardiomyopathy. Persistence is linked to 5’ terminal deletions of viral genomic RNAs that have been detected together with minor populations of full-length genomes in human infections. In this study, we explored the functions and interactions of the different viral RNA forms found in persistently-infected patients and their putative role(s) in pathogenesis. Since enterovirus cardiac pathogenesis is linked to the viral proteinase 2A, we investigated the effect of different terminal genomic RNA deletions on 2A activity. We discovered that 5’ terminal deletions in CVB3 genomic RNAs decreased the proteinase activity of 2A but could not abrogate it. Using newly-generated viral reporters encoding nano-luciferase, we found that 5’ terminal deletions resulted in decreased levels of viral protein and RNA synthesis in singly-transfected cardiomyocyte cultures. Unexpectedly, when full-length and terminally deleted forms were co-transfected into cardiomyocytes, a cooperative interaction was observed, leading to increased viral RNA and protein production. However, when viral infections were carried out in cells harboring 5’ terminally deleted CVB3 RNAs, a decrease in infectious particle production was observed. Our results provide a possible explanation for the necessity of full-length viral genomes during persistent infection, as they would stimulate efficient viral replication compared to that of the deleted genomes alone. To avoid high levels of viral particle production that would trigger cellular immune activation and host cell death, the terminally deleted RNA forms act to limit the production of viral particles, possibly as *trans*-dominant inhibitors.

**Importance:** Enteroviruses like coxsackievirus B3 are able to initiate acute infections of cardiac tissue and, in some cases, to establish a long-term persistent infection that can lead to serious disease sequelae, including dilated cardiomyopathy. Previous studies have demonstrated the presence of 5’ terminally-deleted forms of enterovirus RNAs in heart tissues derived from patients with dilated cardiomyopathy. These deleted RNAs are found in association with very low levels of full-length enterovirus genomic RNAs, an interaction that may facilitate continued persistence while limiting virus particle production. Even in the absence of detectable infectious virus particle production, these deleted viral RNA forms express viral proteinases at levels capable of causing viral pathology. Our studies provide mechanistic insights into how full length and deleted forms of enterovirus RNA cooperate to stimulate viral protein and RNA synthesis without stimulating infectious viral particle production. They also highlight the importance of targeting enteroviral proteinases to inhibit viral replication while at the same time limiting the long-term pathologies they trigger.

## Introduction

Enteroviruses are small non-enveloped viruses from the *Picornaviridae* family. The enterovirus genus includes many viruses associated with human infections: poliovirus, coxsackievirus (CV), enterovirus A71, enterovirus D68, and human rhinovirus. Enterovirus infections result in a wide range of diseases ranging from the common cold to life threatening CNS infections, pancreatitis, or cardiomyopathy [reviewed in(1)]. Like all picornaviruses, enteroviruses are composed of an icosahedral capsid protecting a single-stranded RNA genome of about 7,500 nucleotides (2). Following viral entry, the RNA genome is released in the cytoplasm of the host cell, and translation of the positive-strand RNA results in a unique polyprotein product. The polyprotein is subsequently cleaved by two viral proteinases [2A and 3C (and its precursor 3CD)] to generate the functional proteins required for viral replication and assembly of progeny virions (Fig. 1A) (3). The coding sequences can be delimited into three regions: The P1 region encodes the structural proteins (VP1 to VP4), used for capsid assembly. The P2 and P3 regions encode precursor polypeptides that will be proteolytically cleaved to produce the non-structural proteins required for viral replication, including the RNA-dependent RNA polymerase (3D^pol^). These proteins also mediate the alteration and shut-off of specific host cell functions, primarily as a result of the cleavage activities of the viral proteinases 2A and 3C/3CD (4, 5).

**FIG 1.**
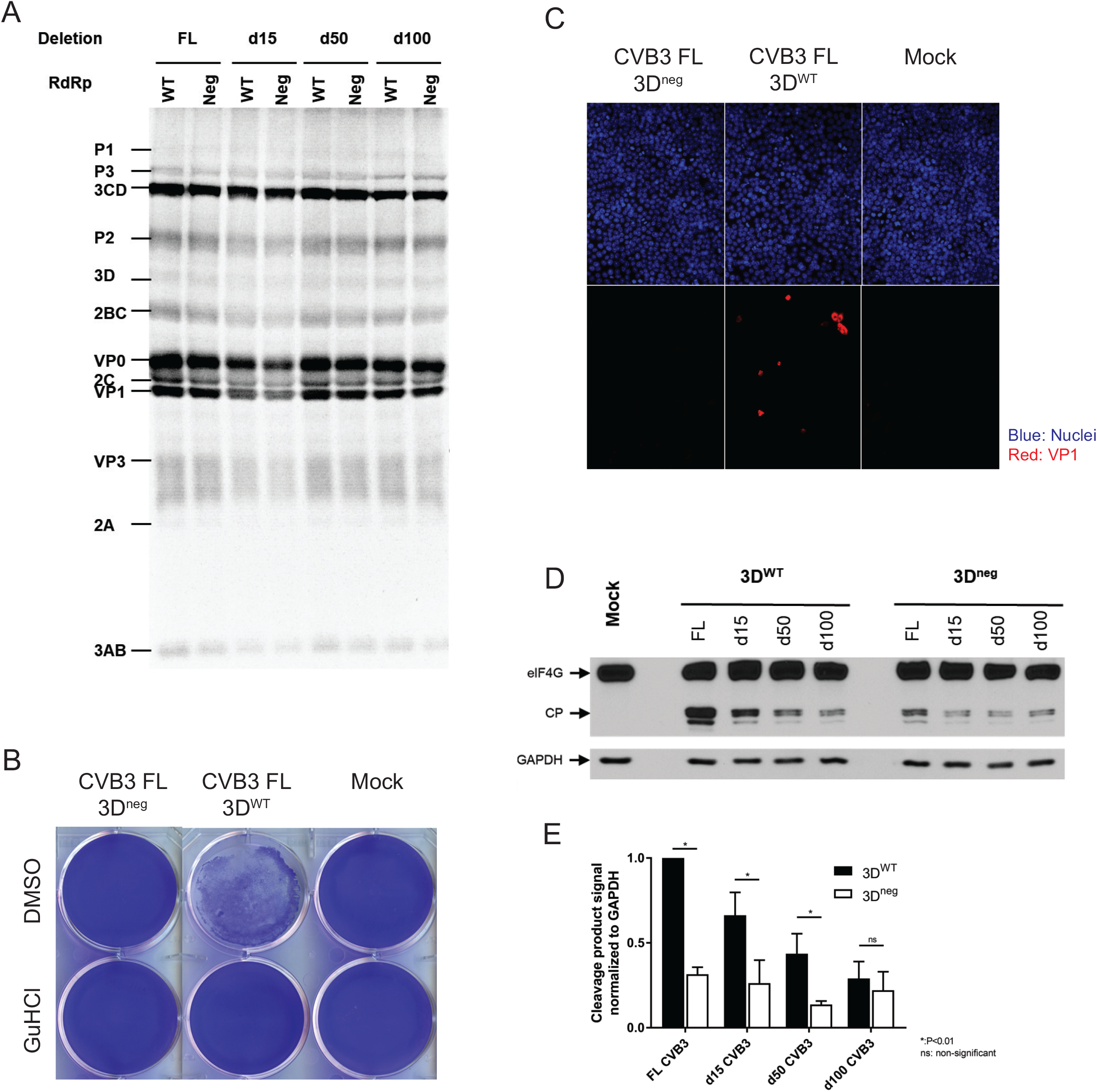
Inactivation of viral 3D polymerase and its effect on 2A proteinase activity. **(A)** *In vitro* translation assay. RNAs were transcribed *in vitro* and incubated for 5.5 hr (4 hr at 30°C followed by 1.5 hr at 34°C) in S10 extract from HeLa cells in the presence of ^35^S methionine. Reaction mixtures were subjected to electrophoresis on an SDS-containing polyacrylamide gel. FL, full-length CVB3 genomic RNA; d15, d50, d100, 5’terminally-deleted CVB3 RNAs harboring deletions of 15 nt, 50 nt, or 100 nt, respectively. RdRp, RNA-dependent RNA polymerase coding sequences in viral RNAs, either wild type (WT) or inactivated by site-directed mutagenesis (Neg). **(B)** HeLa cell viability (determined by crystal violet staining) at 48 hr post-transfection of RNA for CVB3 FL 3D^WT^ or RNA for CVB3 FL 3D^neg^ harboring a polymerase inactivating mutation in the 3D RNA-dependent RNA polymerase coding region in the absence (DMSO) or presence of guanidine hydrochloride (GuHCl). **(C)** Immunofluorescence staining of HeLa cells at 24 hr post transfection of viral RNAs (CVB3 FL 3D^neg^ or 3D^WT^) using antibodies to capsid protein VP1. **(D)** Western blot analysis using antibodies specific for eIF4G. AC16 cardiomyocytes were transfected with viral RNAs [FL or deleted (d15, d50, or d100) forms]. Cells were harvested at 24 hr post-transfection, protein extracts were generated and subjected to SDS PAGE followed by Western blot analysis. Experiments were performed in triplicate and the extent of eIF4G cleavage was quantified using Image J, shown in panel **(E)**.

Enterovirus cardiac infections, especially those caused by coxsackieviruses in group B (CVB), have been reported to result in acute myocarditis that can lead to chronic cardiomyopathies [reviewed in (6)]. The overall prevalence of myocarditis from all causes has been estimated to range from 10 to 100 per 100,000 people worldwide (7). Although numerous etiologies, including other viruses and non-viral pathogens, have been linked to myocarditis, enteroviruses are thought to be responsible for up to 25% of viral myocarditis case (8). Most enterovirus infections are asymptomatic, but those that progress to chronic myocarditis and dilated cardiomyopathy (DCM) can have life-threatening consequences. It has been reported that 10% to 20% of cases of acute myocarditis can progress to a chronic disease state and, ultimately, to DCM and the need for a heart transplant (9, 10). Additional evidence for the involvement of enteroviruses in DCM comes from studies that detected enteroviral capsid protein VP1 as well as genomic RNA in cardiac tissues from patients with late-stage DCM (11, 12). Beyond the use of medication to minimize arrythmia, the implanting of a pacemaker, or heart transplantation, there are no effective anti-viral treatments that would prevent the disease sequelae requiring such dramatic interventions.

The exact mechanisms driving the evolution from acute to persistent cardiac infection are not fully understood. Previous studies revealed that during both acute fulminant myocarditis and dilated cardiomyopathy (DCM), the viral RNA genome undergoes 5’ terminal deletions (10, 13-15). Deletions in patient samples and murine models ranged from 8 to 50 nucleotides (10, 14), while genetically-engineered CVB3 genomic RNAs harboring deletions of up to 78 nucleotides were able to replicate in cell culture, albeit very inefficiently (16). Once these genomic deletions occur, coxsackie B viruses lose their ability to produce detectable cytopathic effects in cell culture, and their replication levels are drastically decreased (17). This defect in replication results in lower levels of both viral genomic RNA and viral protein synthesis, possibly providing a key component that allows for escape from immune surveillance. Bouin and colleagues previously reported the presence of mixed viral populations composed of a small proportion of complete viral genomes (<5%) coexisting with genomic RNAs harboring 5’ terminal deletions in cardiac tissue of chronically infected patients. That study also showed that following transfection of cultured human cardiomyocytes, both viral RNA populations could, independently or in combination, lead to viral proteinase synthesis (10). Enterovirus proteinases, e.g., protein 2A, are known to cleave host cell proteins, resulting in an inhibition and/or alteration of biological processes like cap-dependent protein synthesis and nucleo-cytoplasmic trafficking, leading to the redirection of cellular resources to viral replication [reviewed in (18, 19)]. Of particular significance to cardiac pathologies, CVB3 infection in cardiac cells leads to the disruption of dystrophin, a key protein that links the intracellular cytoskeleton network to transmembrane components of muscle cells, including those in cardiac tissue. This disruption results in increased cell permeability and loss of function, affecting cardiac systole (20). Subsequent studies found that the proteolytic activity of the CVB3 2A proteinase is responsible for this disruption (21). Importantly, dystrophin disruption was linked to disease severity when 50% of cardiac dystrophin was affected (22).

In the present study, we analyzed the activity of the viral 2A proteinase in human cardiomyocytes during CVB3 infections that were initiated with full-length genomic RNAs as well as RNA forms harboring 5’ terminal deletions. We found that 2A proteinase activity could be readily detected in cells transfected with full length or 5’-deleted forms of CVB3 genomic RNAs, even when the viral genome encoded a catalytically inactive form of the viral RNA-dependent RNA polymerase (3D^pol^), rendering these RNAs replication incompetent. Using *in vitro* cultures of human cardiomyocytes and nano-luciferase (NLuc) reporter constructs, we discovered that the full-length and 5’-deleted forms of viral RNA interact to enhance their translation levels when mixed in the proportions observed in patients [19 copies of the deleted form for each full-length RNA; (10)]. As a result, increased levels of viral RNA replication were also observed. However, in the presence of 5’-terminally deleted forms of CVB3 RNAs in cardiomyocyte cells that were subsequently infected with wild-type CVB3, yields of infectious virus were markedly reduced compared to yields produced in the absence of the deleted viral RNAs. Taken together, these results highlight the possible collaboration of full-length viral RNA with RNA forms harboring 5’ terminal deletions, leading to potentiation of CVB3 RNA replication but not lytic virus production. In addition, our studies reveal the high-level activity of the 2A proteinase in cardiomyocytes, even when produced in very low amounts from non-replicating viral RNAs. These data strongly suggest that potential treatments for enterovirus-induced chronic cardiac pathologies should focus on inhibition of viral 2A proteinase functions, which would reduce both the viral replication levels and the cellular destruction directly caused by this key enteroviral enzyme.

## Material and methods

### Cells and viruses

HeLa(S) cells were originally obtained from Eric Stanbridge, University of California, Irvine. Cells were maintained in DMEM (Dulbecco’s Modified Eagle Medium; Gibco, 12800-017) + 0.36% NaHCO_3_ + 1X antibiotic antimycotic solution (Omega Scientific Inc., AA-40) + 10% NCS (Newborn Calf Serum; Omega Scientific Inc., NC-04) at 37°C.

AC16 cardiomyocytes were purchased from Millipore Sigma (ref SCC109). Cells were maintained in DMEM-F12 (1:1) (Gibco, 21041-025) + 12.5% FBS (Fetal Bovine Serum; Omega Scientific Inc., FB-12) + 1X antibiotic antimycotic solution at 37°C.

Plasmids harboring the complete CVB3 genome and the associated 5’ terminal deletions of 15, 50 and 100 nucleotides were kindly provided by Laurent Andreoletti (Université Reims Champagne-Ardenne, Reims, France). Recombinant plasmids encoding a catalytically inactive 3D^pol^ (3D^neg^) were generated using the Q5 Site-Directed Mutagenesis Kit (NEB, E0554S). Nucleotides at position 6891 and 6892 of the CVB3/28 genome were mutated from AT to GC, changing the amino acid from tyrosine to cysteine. A CVB3 genome encoding a NanoLuc luciferase (NLuc) reporter was generated using the In-Fusion HD Cloning Kit (Takara, 638909). NanoLuc luciferase was cloned from the pLenti6.2-Nanoluc-ccdB, a gift from Mikko Taipale (Addgene plasmid # 87078; http://n2t.net/addgene:87078; RRID:Addgene_87078).

### *In vitro* translation of CVB3 RNAs and subsequent polyprotein cleavage

To determine if the viral 3D^pol^ mutation disrupted the translation of 3CD and/or polyprotein processing, an *in vitro* translation assay was carried out by monitoring the incorporation of ^35^S-methionine intro newly synthesized proteins. Briefly, 600 ng of synthetic RNA was added to a 20 µl reaction containing 2 µl of all-4 mix (1 mM ATP, 250 µM CTP, 250 µM GTP, 250 µM UTP, 16 mM HEPES pH7.4, 60 mM KOAc, 30 mM creatine-phosphate, and 400 µg creatine kinase), 1 µl of RNasin (Promega, 40 U/µl), 2 µl of ^35^S-methionine (20 µCi; approx. 1,000 Ci/mmol) and 12 µl of HeLa S10 cytoplasmic extract as previously described (23). The reaction was incubated for 4 hr at 30°C, then for 1.5 hr at 34°C. Laemmli sample buffer (20 µl) was added to the mixture and incubated for 5 min at 100°C. The samples were subjected to electrophoresis on a 12.5% polyacrylamide gel containing SDS at 110 V overnight. The gel was washed 3 times in DMSO for 30 min, once in DMSO-PPO for 30 min, once in deionized water for 45 min and dried. The gel was exposed to a phosphor screen (Molecular Imager FX Imaging screen-K, Bio-Rad) for 24 hr and imaged using the Amersham Typhoon 5.

### Inhibition assay of 3D^pol^ activity

HeLa cells were grown in DMEM (Gibco). They were seeded in 6-well plates (35 mm diameter per well) to be fully confluent at the time of transfection. Before transfection, media was carefully removed and replaced with 2.5 ml of media containing either 0.5% DMSO or 2 mM guanidine hydrochloride (GuHCl), an inhibitor of viral replication, to verify that the 3D^pol^ mutation catalytically inactivated the polymerase. Cells were transfected with 1 µg of CVB3 3D^neg^ or 1 µg of CVB3 WT RNA diluted in 250 µl of Opti-MEM, 5 µl of mRNA Boost Reagent, and 5 µl of TransIT-mRNA Reagent (Mirus Bio). The reaction mixture was incubated for 5 min at room temperature and added to the cells. Plates were incubated at 37°C with 5% CO_2_ and media was collected after 8 hr.

### Quantification of lytic viral particle production

Plaque assays were used to determine infectious viral yields of CVB3. Six-well plates were seeded with HeLa cells to be fully confluent at the time of staining. Media was removed, and cells were washed twice with PBS. Adsorption was carried using 200 µl of collected media from previously transfected cells and incubated for 30 min at room temperature. After incubation, an overlay of 0.45% agarose containing DMEM supplemented with 10% NCS, 10 mM MgCl_2_ and 20 mM HEPES was added, and plates were incubated at 37°C with 5% CO_2_ for 2 to 3 days for CVB3 or CVB3 encoding NanoLuc luciferase (NLuc), respectively. Cells were fixed with 10% TCA and overlays were removed. Cells were stained with crystal violet and washed with water to visualize virus plaques.

### Immunofluorescence assays

HeLa cells were seeded onto glass coverslips in a 24-well plate to be 70-80% confluent at the time of transfection with 0.5 ml of media. Cells were transfected with a total of 1 µg of viral RNA diluted in 50 µl of Opti-MEM, 1 µl of mRNA Boost Reagent and 1 µl of TransIT-mRNA Reagent (Mirus Bio) and incubated for 5 min at room temperature. The mixture was added to the cells, and the plates were incubated at 37°C with 5% CO_2_ for 24 hr. After incubation, media was removed and cells were fixed and permeabilized with a cold methanol-acetone mixture (3:1) and incubated for 10 min at -20°C. Cells were washed 3 times with phosphate-buffered saline (PBS) and blocked in PBS supplemented with 1% BSA and 2% FBS for 30 min at room temperature prior to primary antibody incubation. Cells were incubated with anti-VP1 antibodies (Dako, anti-enterovirus clone5-D8/1) diluted 1:500 in blocking solution for detection of viral protein VP1 or anti dsRNA antibody (SCICONS, anti-dsRNA mAb J2, 10010500) diluted 1:1000 in blocking solution overnight at 4°C. Cells were washed 5 times with PBS before being incubated with goat anti-mouse IgG heavy and light chain Dylight-conjugated secondary antibodies (Bethyl) diluted 1:400 in blocking solution for 1 hr at 37°C followed by 5 washes with PBS. Cells were incubated with DAPI diluted 1:250 in PBS for 2 min at room temperature followed by washing once in PBS. Coverslips were mounted on a slide with Fluoro-Gel (Electron Microscopy Sciences) and imaged.

### 2A proteinase activity assay using cleavage of eIF4G

The activity of the viral 2A proteinase was assessed using a Western blot assay targeting eIF4G. AC16 were seeded in 6-wells plate to be 80% confluent on the day of transfection. Cells were transfected with 2.5 µg of CVB3 genomic RNA diluted in 250 µl of Opti-MEM, 5 µl of mRNA Boost Reagent, and 5 µl of TransIT-mRNA Reagent (Mirus Bio). The reaction mixture was incubated for 5 min at room temperature and added to cell monolayers. Plates were incubated at 37°C with 5% CO_2_ for 24 hr. Cells were lysed using RIPA buffer and total protein concentration was quantified using Bio-Rad Protein Assay Dye Reagent Concentrate (Bio-Rad, 500-0006). Twenty micrograms of protein were incubated in Laemmli sample buffer at 95°C for 5 min and subjected to electrophoresis on a 3-15% gradient polyacrylamide gel. Proteins were transferred to a PVDF membrane (Immobilon-P, Millipore, IPVH00010) at 100 mA overnight. The resulting membrane was blocked for 1 hr in PBS + 5% BSA (Fisher Bioreagents, BP1605-100) before being incubated overnight with primary antibodies [anti-VP1 (Dako, anti-enterovirus clone5-D8/1) or anti-eIF4G (Cell Signaling Technology, 2469S) diluting in blocking buffer at 1:500 and 1:1000, respectively]. Membranes were washed 5 times in TBS + 0.1% Tween 20 (Sigma Aldrich, P1379) before being incubated for 1 h at room temperature with secondary antibodies HRP conjugated (Bethyl, A90116P or A120-101P) diluted 1:20 000 in TBS + 0.1% Tween 20. Membranes were treated with ECL (Thermo Scientific, 32106) and imaged using an Amersham Imager 680.

### Quantification of viral protein synthesis using a bioluminescent reporter

NanoLuc luciferase (NLuc) was cloned into the CVB3 genome upstream of the P1 coding sequence. Quantification of viral protein expression was performed by monitoring the bioluminescent signal produced by NLuc. AC16 cells were transfected with indicated quantities of viral RNA and incubated for 24 hr. Cell monolayers were then scraped and subjected to 3 freeze/thaw (dry ice/37°C) cycles. Samples were diluted and the bioluminescent signal was quantified using Nano-glo Luciferase Assay (Promega, N1110). Readings were performed on a Berthold Sirius luminometer.

## Results

### 2A proteinase activity following cardiomyocyte transfection of replication-deficient CVB3 RNA

Bouin and colleagues previously reported that viral forms harboring terminal deletions exhibited 2A proteinase activity when transfected into primary human cardiomyocytes, even with undetectable levels of viral capsid protein (10). To expand our understanding of the effect of active replication on 2A proteinase activity, we generated CVB3 genomic RNA transcripts with 5’ terminal deletions of 0, 15, 50, or 100 nucleotides (nts) that also encoded an inactive 3D polymerase (3D^neg^) (Fig. 1). We analyzed polyprotein processing following *in vitro* translation in HeLa cell S10 extracts and did not detect any defects in the production of mature CVB3 proteins (Fig. 1A). Viral RNA was then transfected into HeLa cell monolayers to assay for cytopathic effects (CPE) and viral protein production (Fig. 1B and 1C, respectively). Cytopathic effects were assessed following crystal violet staining 48 hr post transfection, with or without addition of guanidine hydrochloride (GuHCl) to inhibit viral RNA replication. Complete CPE was observed in cells transfected with wild-type CVB3 RNA and was completely inhibited by GuHCl. In addition, no CPE were observed following transfection of CVB3 3D^neg^ genomic RNA, which harbors a catalytic site mutation that ablates RNA synthesis activity (Fig. 1B). Similarly, immunofluorescence staining to detect viral protein VP1 did not reveal any viral protein production in CVB3 3D^neg^-transfected cells compared to those transfected with wild-type CVB3 RNA. These results validated our CVB3 3D^neg^ construct as a defective polymerase-encoding RNA.

To determine the effects of a non-replicating RNA on 2A proteinase activity in cardiac cells, we performed transfections of the AC16 cardiomyocyte cell line. At 24 hr post-transfection, cells were harvested and extracted proteins were subjected to Western blot analysis to detect eIF4G (Fig. 1D). All viral RNAs tested (including those with 5’ terminal deletions) displayed 2A proteinase activity, as eIF4G cleavage was observed, regardless of 3D polymerase functionality. Since the CVB3 genome is a positive-strand RNA, the transfected genome can be used as a template to produce 2A proteinase in the absence of RNA replication and template amplification. The extent of cleavage of eIF4G is progressively reduced as the size of the 5’ terminal deletions increases, and these levels are reduced further in the absence of RNA replication, as observed in Fig. 1D. However, the readily detectable signal that we observed even for the 100 nt deletion construct in the absence of RNA synthesis provides compelling evidence that 2A proteinase activity is very high in these transfected cardiomyocytes suggesting, in part, how viruses harboring terminal deletions can still induce cardiac pathology. These experiments were performed in triplicate, and the cleavage products were quantified. The results are presented in Fig. 1E. As expected, the longer the deletion, the lower 2A proteinase activity was observed. Interestingly all deletions observed in patients (up to 50 nts) had a statistically significant decrease in 2A activity when 3D polymerase was inactivated. The deletion of 100 nucleotides removes the entire stem loop I RNA structure, eliminating the binding site for the proteinase-polymerase precursor polypeptide 3CD that is required for viral RNA replication (24, 25); it is therefore not surprising that no differences in 2A proteinase activity were observed between the RNAs encoding either an active or inactive form of the 3D. These results provide evidence that viral RNAs with 5’ terminal deletions up to 50 nucleotides can replicate and amplify their RNA in cells after transfection, even though they produce levels of viral proteins that are not detectable by Western blot analysis [refer to Fig. 5 in (10)].

### Generation of recombinant forms of CVB3 expressing nano-luciferase

Monitoring eIF4G cleavage fragments resulting from 2A proteinase activity was insightful for our analysis, but this approach provided only indirect proof of replication of the 5’ terminally-deleted forms of CVB3 RNAs. To implement a more specific reporter of viral replication and protein synthesis, we generated a recombinant form of CVB3 that encodes nano-luciferase (NLuc) (Fig. 2A). The NLuc gene was inserted on the 5’ extremity of the polyprotein coding sequence, followed by a 3C/3CD cleavage site to ensure proper processing of the viral polyprotein. Viral genomic RNA was synthesized *in vitro* and transfected into HeLa cells. Recovered NLuc reporter virus was able to undergo a complete replication cycle and induced total CPE within 24 hr, similar to that produced by WT CVB3 (Fig. 2B). Quantification of infectious particle production was performed by plaque assay. Viruses expressing NLuc exhibited a decrease in viral titer of ∼90% at both 8 hr and 24 hr post-infection (Fig. 2C). Interestingly, this defect did not result in reduced levels of RNA synthesis during the first 8 hr of infection, which is the normal time for WT CVB3 to complete one replication cycle in cultured HeLa cells (Fig. 2D). An *in vitro* translation assay was performed on transcripts corresponding to WT CVB3 genomic RNA and compared to genomic RNA transcripts expressing NLuc and harboring deletions of 0, 15, and 50 nts. Production of the non-structural proteins (e.g., 3CD) was similar between WT and the different 5’ terminally-deleted forms of RNA encoding NLuc (Fig. 2E). However, polyprotein processing of the capsid precursor (P1) was incomplete, resulting in an accumulation of the NLuc-P1 precursor and reduced levels of mature capsid proteins (e.g., VP0 and VP3). The reduced levels of capsid proteins produced in cells infected with the CVB3 NLuc virus could account for the decrease in viral titers reported in Fig. 2C, possibly due to an inefficient encapsidation step. This result would also explain the near WT levels of RNA synthesis by the NLuc recombinant virus shown in Fig. 2D, since impaired production of mature capsid proteins would not be expected to reduce intracellular levels of viral RNA. Taken together, these results validate the NLuc recombinant virus as a reporter for viral RNA replication and protein synthesis.

**FIG 2.**
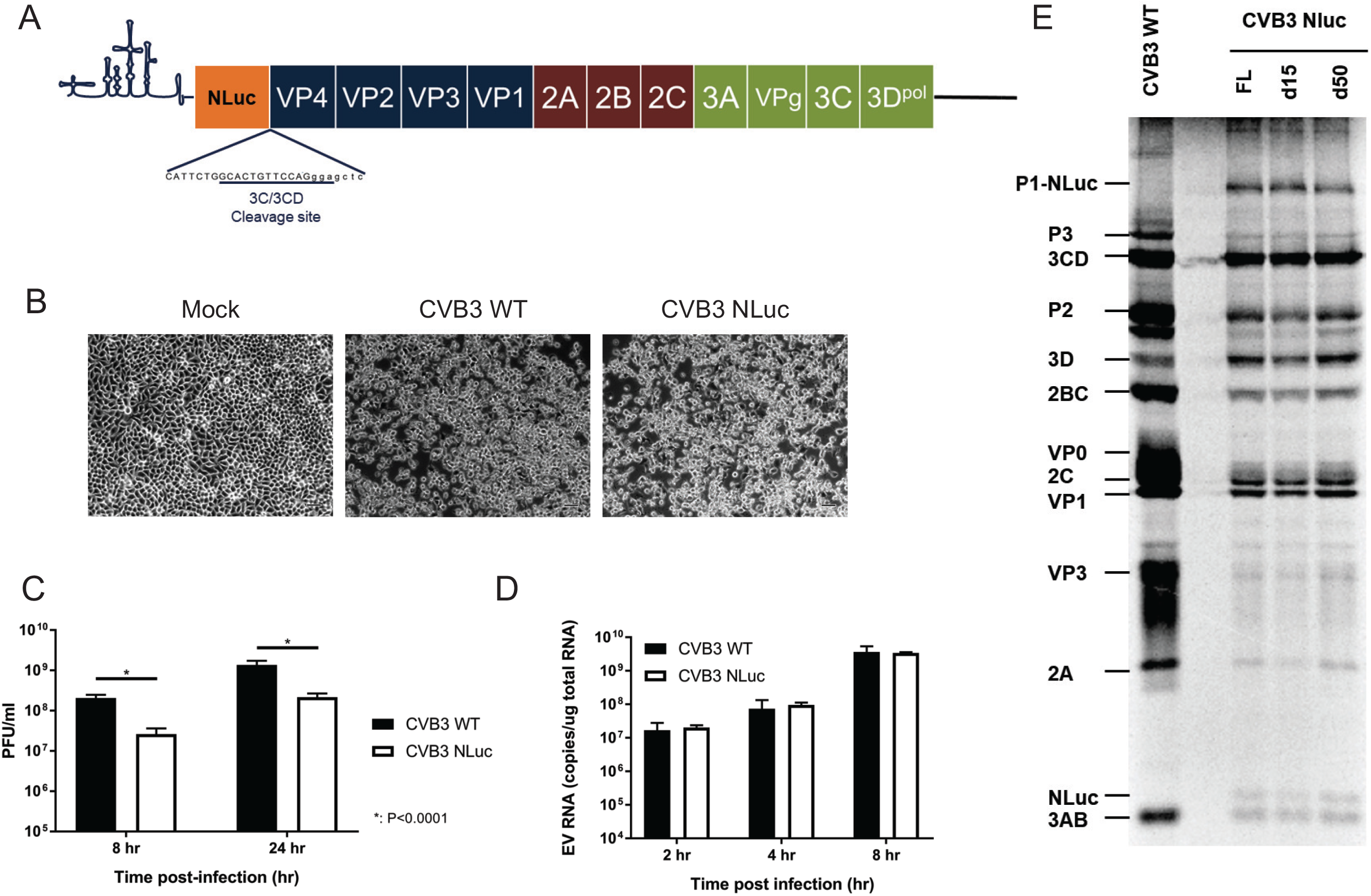
Generation and characterization of CVB3 recombinant genomes expressing nano-luciferase. **(A)** Schematic representation of CVB3 genome encoding NanoLuc luciferase (NLuc). *Virus stocks were generated in HeLa cells following transfection with WT viral RNA or* CVB3 RNA expressing nano-luciferase. **(B)** CPE in HeLa cells was observed at 24 hr post-infection for both CVB3 WT- and CVB3 NLuc-infected cells but not mock-infected cells. **(C)** Infectious viral particle production in HeLa cells at 8 hr or 24 hr post-infection was quantified by plaque assay. **(D)** Viral RNA quantification in cells infected with CVB3 WT or CVB3 NLuc at 2 hr, 4 hr, or 8 hr post-infection was performed by RT-qPCR. **(E)** *In vitro* translation assay. RNAs were transcribed *in vitro* and incubated in S10 extract from suspension HeLa cells in the presence of ^35^S methionine. Reaction mixtures were subjected to electrophoresis on an SDS-containing polyacrylamide gel followed by fluorography and phosphorimaging analysis.

### Viral protein synthesis directed by 5’ terminally deleted CVB3 genomes

Using viral nano-luciferase reporters, we analyzed the protein synthesis capacities of viral genomic RNAs harboring 5’ terminal deletions. *In vitro* transcribed RNAs were incubated in S10 extracts derived from suspension HeLa cells. Nano-luciferase substrate was added to the reaction and luminescence was quantified (Fig. 3A). As expected, genomes without deletions (FL) produced the highest levels of viral proteins. Viral genomes harboring terminal deletions of 15 and 50 nucleotides had a signal decrease by 80% and 40%, respectively. Although somewhat counterintuitive to expectations, the larger deletion of 50 nts had less of a negative impact on viral translation compared to the small deletion of 15 nts. Since neither deletion extends into the IRES sequences required for CVB3 translation, it is possible that the smaller deletion disrupts the overall folding of the RNA to a greater extent compared to the larger one.

**FIG 3.**
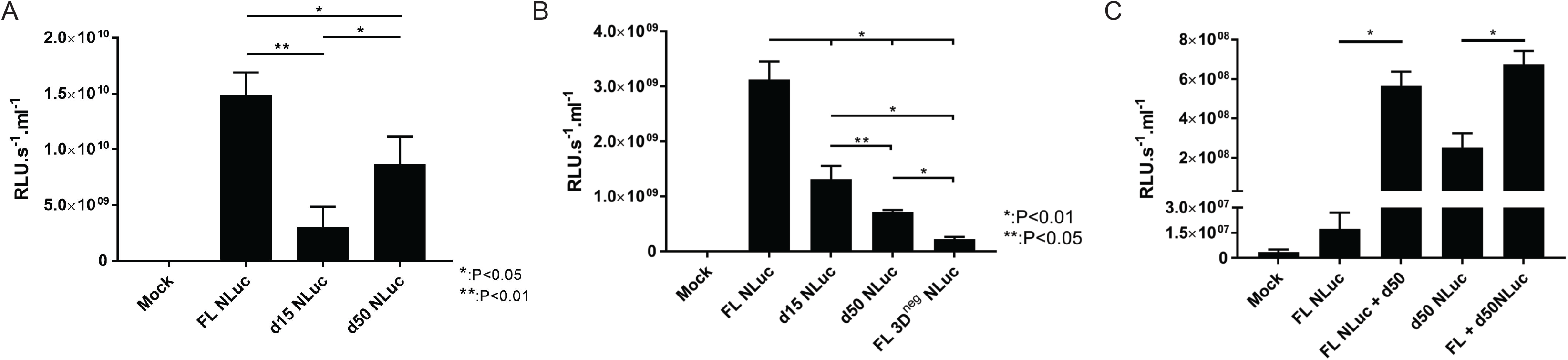
Viral protein synthesis by 5’ deleted and full length CVB3 RNAs in cell culture. **(A)** *In vitro* translation assay of 5 ‘deleted and full length CVB3 genomic RNAs. RNAs were transcribed *in vitro* and incubated in S10 extracts from HeLa cells for 4 hr at 30°C. NanoLuc luciferase substrate was added, and luminescence was quantified. **(B)** AC16 cardiomyocytes were transfected with viral RNAs encoding nanoLuc luciferase (NLuc) and harboring 5’ terminal deletions. Cells were lysed after 24 hr by freeze/thaw cycles, NLuc substrate was added, and luminescence was quantified. **(C)** AC16 cardiomyocytes were transfected with a mixture of deleted viral RNA (d15 or d50) and full length (FL) RNA in a 19:1 ratio. To identify the template driving viral protein synthesis, only one RNA form encoded NLuc. Cells were lysed after 24 hr by freeze/thaw cycles, NLuc substrate was added, and luminescence was quantified

The *in vitro* translation assay used to produce the data shown in Fig. 3A does not recapitulate the RNA synthesis portion of the viral cycle. To overcome this limitation, RNAs were transfected into AC16 cardiomyocyte cells (Fig. 3B). As expected, RNAs harboring 5’ terminal deletions produced a weaker signal than full-length genomes, and more than genomes harboring an inactive 3D polymerase (3D^neg^). Interestingly, RNAs harboring deletions of 15 nts produced a stronger signal compared to those harboring deletions of 50 nts. That difference between the *in vitro* and the cell culture assays might be explained by the RNA synthesis capacities of the viral RNAs in the AC16 transfection model that cannot be recapitulated *in vitro*, since these latter assay conditions do not permit viral RNA replication.

To study the interactions between the full length (FL) and deleted (d50) forms of CVB3 genomic RNAs found in persistently-infected cardiac tissues, we co-transfected AC16 cells in the same proportion as detected in patients in previous reports (10), i.e., nineteen 5’ terminally deleted genomic RNAs for each full-length genomic RNA [19:1] ratio (Fig. 3C). To discriminate between proteins produced by the two different RNA forms following co-transfections, only one form encoded the NLuc reporter. Interestingly, when co-transfected, both forms produced a higher luminescent signal than they produced individually, with a 30-fold increase when the reporter was encoded on the FL RNA and 2.5-fold increase when the reporter was encoded by the d50 form. Note that for the singly transfected samples Fig. 3C (FL NLuc versus d50 NLuc), the input amount of FL RNA was 1/19^th^ of that used for the d50 NLuc sample, thus accounting for the apparent discrepancy in these data. This experiment provides the first evidence of synergistic interactions between the different persistent forms of enterovirus genomic RNAs in cell culture.

### RNA synthesis directed by CVB3 genomic RNA transfected into cardiomyocytes

During viral replication, protein and RNA synthesis are intertwined and inter-dependent. Since the data shown in Fig. 3 demonstrated a stimulation of viral protein synthesis when the different viral RNA forms were co-transfected (as measured by NLuc reporter signals), we investigated the effects of the 5’ terminal deletions on viral RNA synthesis. Viral RNAs were transfected into AC16 cardiomyocytes, and viral RNA was quantified by RT-qPCR targeting 5’UTR (Fig. 4A). RNA harboring deletions of 15 nucleotides did not have a statistically different RNA signal compared to full-length forms, but a trend showed a slight decrease in mean values of 12.6%. However, viral forms with 50 nt deletions showed a clear decrease of ∼33%. This difference could explain the discrepancy between the protein synthesis levels observed *in vitro* compared to those obtained following transfections of cultured AC16 cells. The defect in translation for the 15 nt deleted form can be rescued by increased levels of viral RNA synthesis.

**FIG 4.**
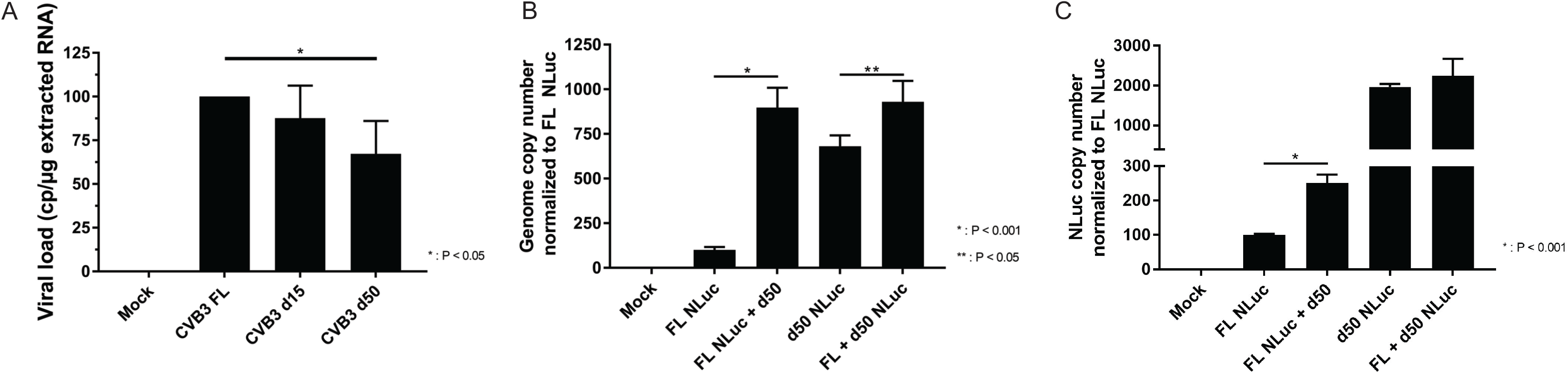
Viral RNA synthesis by 5’ deleted and full length CVB3 RNAs in in cell culture. **(A)** AC16 cardiomyocytes were transfected with viral RNAs encoding NLuc and harboring 5’ terminal deletions. RNA was extracted and RT-qPCR targeting the 5’UTR was performed. **(B)** AC16 cardiomyocytes were transfected with mixtures of deleted viral RNA and FL RNA in a 19:1 ratio. To determine the origin of the template driving viral RNA synthesis, only one RNA form encoded NLuc. RNA was extracted and RT-qPCR targeting the 5’UTR was performed. **(C)** As in (B), except NLuc sequences were targeted for RT-qPCR reactions. Results in (B) and (C) are normalized to the average value of FL NLuc transfection.

To confirm the findings on the interactions between the 5’ deleted and full-length RNA forms, we performed co-transfection assays in AC16 cells in the [19:1] ratio described previously. RNA accumulation was assessed by RT-qPCR using primers specific for 5’ UTR sequences (Fig. 4B) to quantify total viral replication and for NLuc (Fig. 4C) sequences to discriminate which RNA form had increased levels of RNA replication. Measurements using 5’ UTR sequence amplification highlighted an increase in viral RNA when both forms were mixed (Fig. 4B), confirming the interaction described above. Results from the NLuc-specific amplification showed an increase in full length RNA when mixed with forms harboring 5’ terminal deletions; however, we did not observe statistically significant differences for the RNA forms harboring 50 nt terminal deletions. These results suggest that despite both RNA forms benefitting from interactions following co-transfection, the mechanism leading to the viral protein synthesis increase appears to be indirect. Full length forms are producing more RNA, which results in a higher number of viral templates for translation. Deleted forms are not efficient templates for RNA replication; however, they may benefit from the activity of proteins synthesized following the large amplification of full-length RNAs during RNA replication that redirect cell resources to viral translation.

To track active RNA synthesis *in situ*, we performed RNA transfections in HeLa cells followed by immunofluorescent staining for double stranded RNA (dsRNA) (Fig. 5). This RNA form is produced during viral RNA replication, primarily during negative-strand RNA synthesis occurring on replication organelles. As expected from previous results, full-length genomic RNA (FL) samples produced high levels of signal, while the signal produced by the 5’ deleted RNA (d50) was barely detectible. When both forms were mixed and transfected into HeLa cells, an increase in dsRNA signal was observed. These results confirm what we had observed in previous experiments.

**FIG 5.**
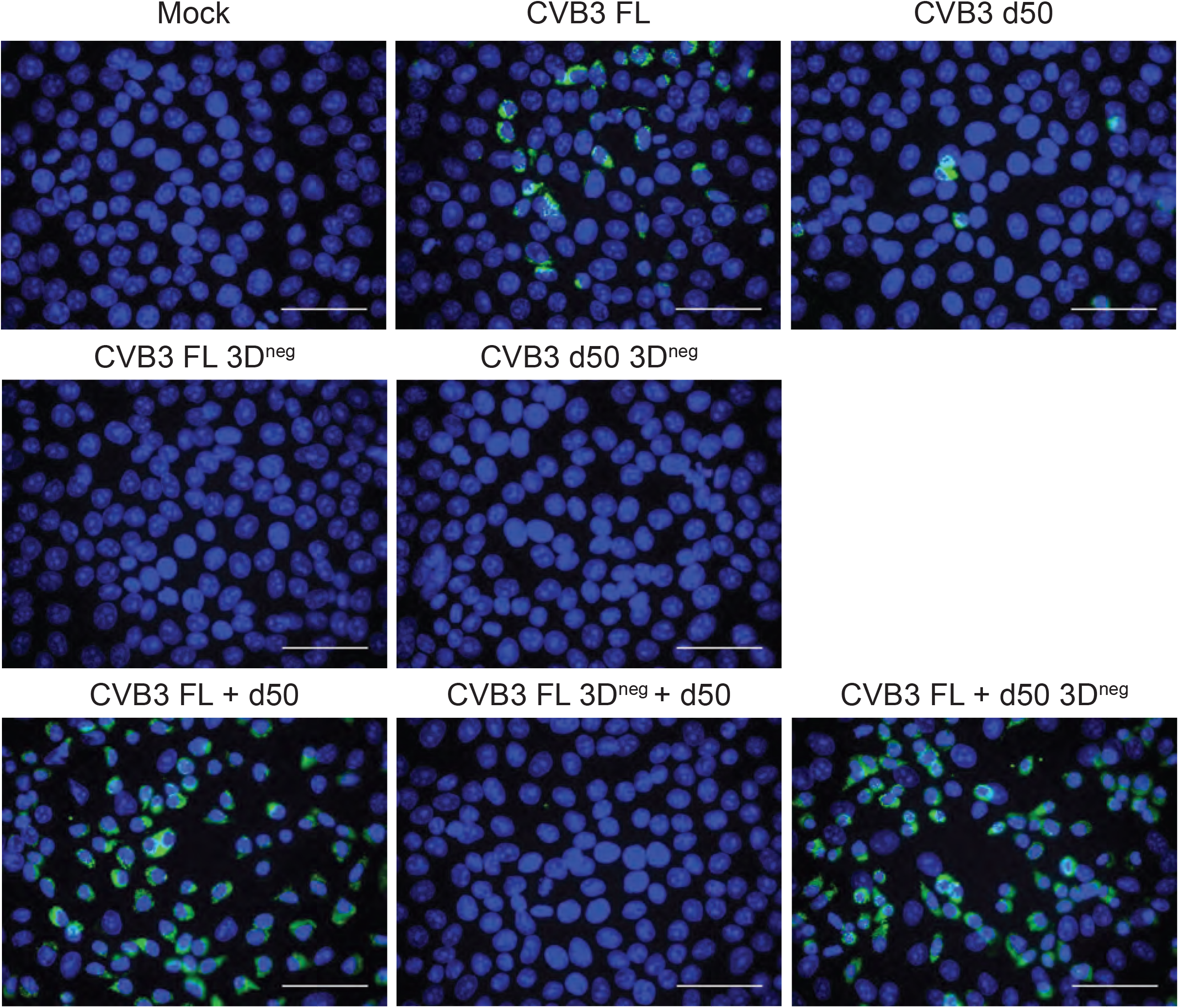
CVB3 RNA synthesis and formation of dsRNA during infection of cultured cells. HeLa cells were transfected with different forms of viral RNA. When two forms of viral RNAs [d50 (5’ deleted) and FL (full length)] were co-transfected, the ratio was 19:1 of 5’ terminally deleted form to FL form, respectively. At 24 hr post transfection, cells were fixed and permeabilized, and immune staining with antibodies specific for dsRNA was performed. Nuclei were stained in blue, and dsRNA in green. 3D^neg^ denotes viral RNAs harboring a genetically inactivated 3D RNA-dependent RNA polymerase. White bar = 50 µm.

To further investigate the role of each RNA form (5’ deleted and full length), we used RNA constructs encoding a catalytically inactive 3D RNA polymerase. As expected, both 5’ terminally and FL forms harboring an inactive 3D^pol^ were not able to produce dsRNA when transfected independently. When full length RNA encoding an inactive 3D polymerase was co-transfected with d50 RNA harboring an active 3D polymerase, no signal was detected. This result is in agreement with the data shown in Fig. 4C, confirming that the full-length template RNAs are the main drivers of RNA synthesis. Co-transfecting full-length genomic RNAs encoding an active 3D polymerase with deleted RNAs encoding an inactive 3D polymerase did not decrease the signal observed for dsRNA staining compared to co-transfection of forms both encoding active polymerases. This result suggests that the defective genome did not act as a dominant-negative for viral RNA synthesis. These results demonstrate that both WT and deleted forms can form replication organelles and carry out viral RNA synthesis in cardiomyocytes. During their interaction resulting from co-transfection, the full-length RNA forms appear to have enhanced RNA and protein synthesis, while the 5’ terminally deleted forms only have an increased protein production.

### Effect of 5’ terminally-deleted CVB3 genomes on new infections

Following our previous results from co-transfections of 5’ terminally-deleted genomic RNA with full-length CVB3 RNAs, we investigated if the presence of preexisting viral RNA harboring 5’ terminal deletions in host cells had any effect on the production of WT CVB3 infectious particles. We transfected HeLa cells or AC16 cells with viral RNAs harboring 5’ terminal deletions of 50 nts. At 24 hr post-transfection, we infected with WT CVB3 or CVB3 encoding NLuc. At 24 hr post-infection, cell monolayers were harvested, and the production of infectious virus particles was quantified by plaque assay. Dose-dependent inhibition was observed in HeLa cells pre-transfected with increasing amounts of 5’ terminally-deleted CVB3 RNA (d50) (Fig. 6A). No inhibition was detected following transfection of 500 ng of RNA. A non-statistically significant decrease of 8% was observed following transfection of 1 µg of d50 RNA, while a significant drop to 40% of control levels was observed following transfection of 2.5 µg of d50 RNA. Similar results were observed for transfection/infection of AC16 cardiomyocytes (Fig. 6B), with a more dramatic inhibition of 90% of virus titers following transfection of 2.5 µg of d50 RNA. Remarkably, this drop in lytic viral particle production did not correspond to a decrease in viral protein levels, as the levels of NLuc signal were nearly identical for d50 transfected AC16 cells versus mock-transfected cells (Fig. 6C). These unexpected results may explain how enteroviruses can persist in cardiac tissue by reducing the levels of lytic particles produced, possibly by inhibiting capsid production or another step in particle assembly/genome packaging.

**FIG 6.**
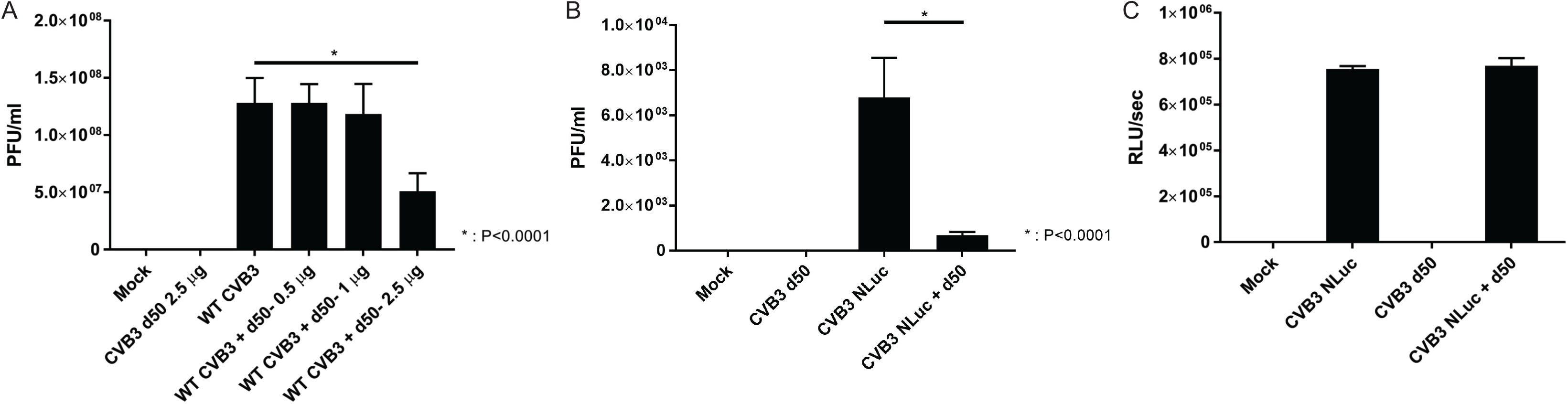
Effect of 5’ terminally-deleted CVB3 genomes on new infections. **(A)** HeLa cells were transfected with the indicated viral RNA quantities harboring a 50 nt 5’ terminal deletion (d50). At 24 hr post-transfection, monolayers were infected with WT CVB3 at a multiplicity of infection (MOI) of 1. At 24 hr post-infection, lytic viral particle production was quantified by plaque assay. **(B)** AC16 cells were transfected with 2.5 µg of viral RNA harboring a 50 nt 5’ terminal deletion. At 24 hr post-transfection, monolayers were infected with Nluc-CVB3 (MOI of 1). At 24 hr post-infection, lytic viral particle production was quantified by plaque assay. **(C)** Viral protein production was assayed by measuring the bioluminescent signal produced by NLuc in the experiment described in (B).

## Discussion

Previous studies on persistent enterovirus infections during chronic cardiac pathologies revealed that viral replication is decreased and viral particle production is dramatically reduced (6, 26, 27). During the progression from acute to persistent cardiac infection (outlined schematically in Fig. 7), the enterovirus genomic RNA undergoes deletion of some of its 5’ terminal nucleotides (28). Cardiac samples originating from patients suffering from long-term dilated cardiomyopathy were analyzed, and this analysis revealed that these terminal deletions can range up to 50 nucleotides, impacting the binding sites of host cell protein PCBP2 and viral protein 3CD/3D on stem-loop I of the viral genomic RNA (10, 29). While it was shown that full-length and deleted genomic RNA forms were both able to produce 2A proteinase activity *in vitro* and in transfected cardiomyocytes (10, 29), an important factor in DCM pathogenesis, the exact role of each RNA form in RNA replication and protein synthesis remained unknown. In the current study, we aimed to further investigate the 2A activity of these different forms and its impact on host functions.

**FIG 7.**
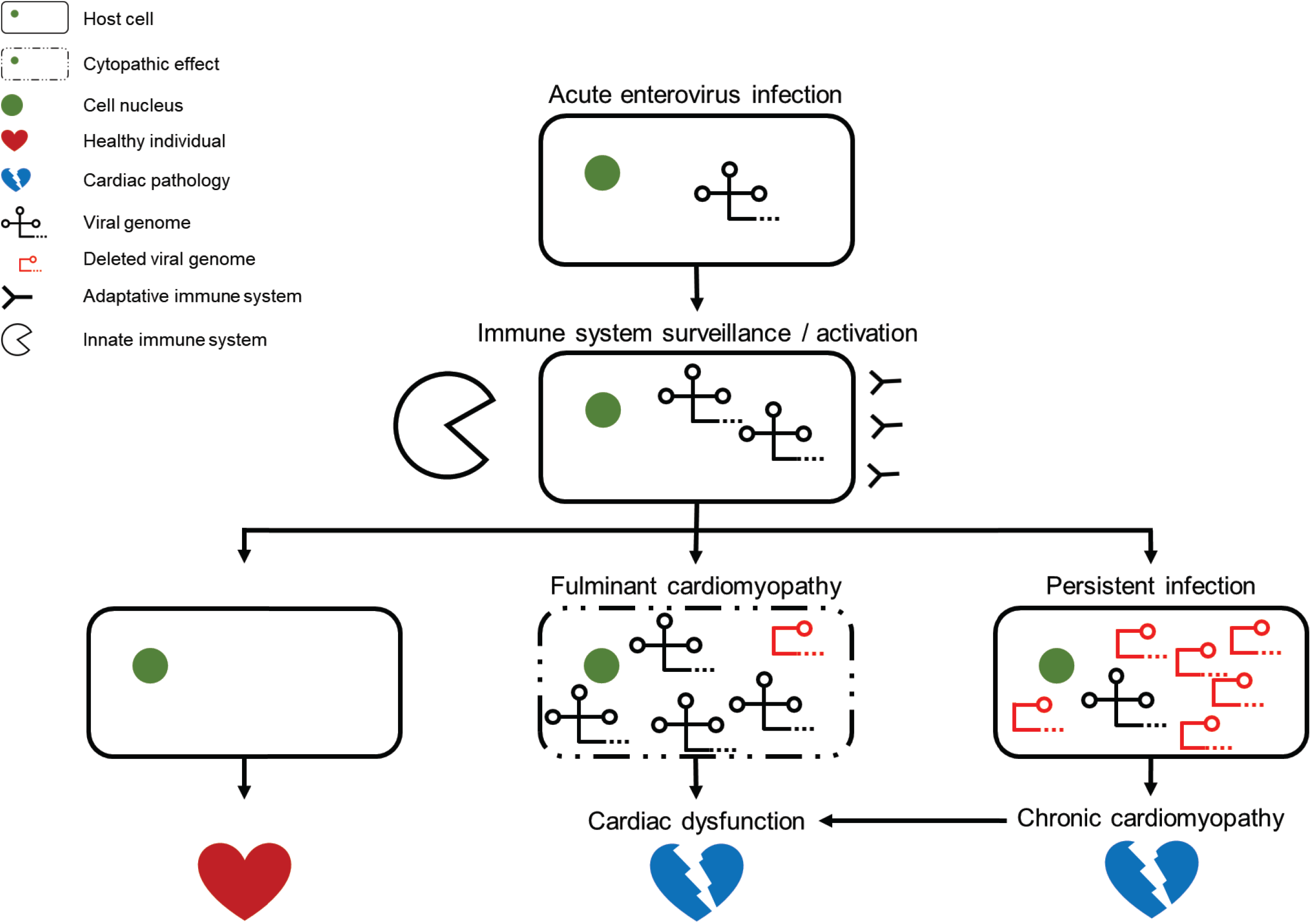
Schematic representation of possible enterovirus cardiac infection outcomes. Acute infection by wild-type enterovirus leads to the activation of the host immune response. Depending on the individual, the infection can be shut down by the immune defense, resulting in viral clearance without any major sequelae for the host. Alternatively, the infection can induce a fulminant cardiomyopathy, inducing high levels of cell pathology, leading to severe, life-threatening, cardiac dysfunction. During fulminant cardiomyopathy, both full-length and 5’ terminally deleted forms of genomic RNA have been observed in infected cells. In some case, the immune system will help to clear most of the acutely infected cells, but some cells will remain persistently infected. To escape from the host immune surveillance, persistent viral populations will mainly be comprised of RNA forms that harbor 5’ terminal deletions with minor full-length RNA forms and will exhibit very low levels of virus replication. The interactions of these different RNA forms lead to viral protein and RNA production while limiting the production of infectious viral particles. The persistent infection will ultimately lead to a chronic cardiomyopathy, evolving slowly to cardiac dysfunction and the need for a heart transplant.

Since viral protein expression in human cardiomyocytes transfected with 5’ terminally deleted genomes could not be detected by Western blot assay (10), the first question we addressed was whether active replication was needed to trigger detectable 2A proteinase activity and subsequent DCM pathogenesis. We generated cDNA clones harboring 5’ terminal deletions of 0, 15, 50 and 100 nucleotides, expressing either an active wild-type 3D RNA-dependent RNA polymerase or a mutated form encoding a catalytically inactive version of this viral protein. We confirmed that the inactivating mutation did not impair the proteinase activity of the 3CD precursor of the polymerase by analyzing polyprotein processing using an *in vitro* translation assay (Fig. 1A). In a second step, we confirmed that replication was abolished by monitoring cytopathic effects (Fig. 1B) and the synthesis of viral capsid protein VP1 (Fig. 1C). To assess 2A activity in our different genomic constructs, we transfected viral RNAs into AC16 cardiomyocytes and performed Western blot assays using antibodies to eIF4G. eIF4G is a well-known target of 2A, and its cleavage is part of the viral mechanism to shut down and redirect cell host translation mechanism to support viral replication. As expected, increasing the size of 5’ terminal deletions correlated with a decreasing 2A proteinase activity. Non-deleted genomic RNAs (WT) produced the highest level of 2A-dependent cleavage of eIF4G. RNA harboring 5’ terminal deletions of 15 and 50 nucleotides, mimicking populations detected in patients suffering from chronic cardiac pathologies, showed a decrease in eIF4G cleavage of 34% and 56%, respectively. Finally, the genomic RNA engineered to harbor a 5’ terminal deletion of 100 nucleotides, removing the entire stem-loop I and, more specifically, the 3CD protein binding site, had a decrease in 2A proteinase activity of 71%. Interestingly, all of the CVB3 RNAs encoding a catalytically inactive 3D polymerase had similar activity, with a decrease of 70% or more compared to the WT full-length form. These results underscore the very high level of catalytic activity of the 2A proteinase and confirm that active replication is not required to produce quantities of this enzyme still capable of inducing detectable proteolytic activity leading to intracellular pathology. Since the enterovirus genomes are positive strand polarity, a small amount of RNA is enough to produce 2A. It is important to note that our experiment was carried out over a 24 hr period, as opposed to our previously described patient samples where RNA was analyzed in cardiac tissue (10). The length of time between diagnosis and cardiac sampling was, on average, 105 months, leaving an extended period of time for 2A activity even with a very low level of viral replication. Collectively, these results highlight the ability of viral RNA forms harboring 5’ terminal deletions to produce 2A proteinase activity.

In a previous study (10), Bouin and colleagues demonstrated that deleted forms of enterovirus RNAs account for the major proportion of the viral population in cardiac tissues of human patients with idiopathic DCM; however, these RNAs are almost always associated with a low level (1-5% of total viral population) of full-length genomic RNAs; refer to Fig. 7. The next set of experiments was carried out to understand the dynamics and consequences of these interactions. To monitor the different RNA forms independently, we engineered viral RNA constructs with a bioluminescent reporter. As 2A activity is directly dependent on viral translation efficiency, we engineered NanoLuc luciferase (NLuc) into RNAs harboring different 5’ terminal deletions and validated the biological activities of these constructs (Fig. 2). Viral protein synthesis activity was assessed for each of the constructs by *in vitro* translation assays (Fig. 3A) and following transfections of AC16 cardiomyocytes (Fig. 3B). Interestingly, a discrepancy was observed for RNA forms harboring 5’ terminal deletions of 15 or 50 nucleotides. *In vitro* experiments showed a higher level of CVB3 protein synthesis for the 15 nucleotide deletion (d15) construct [compared to the 50 nucleotide deletion (d50)], but RNA transfection assays in cultured AC16 cells produced the opposite result. This difference can be explained by the different RNA synthesis activity of the different forms (Fig. 4A). Our *in vitro* translation assay does not recapitulate the full replication cycle, as RNA synthesis is very inefficient in this model. The defect in RNA synthesis observed for the d50 form in transfected cells would not be relevant in the cell-free translation assay, which allows quantification of viral protein synthesis levels uncoupled from the RNA synthesis that occurs following transfection. The *in vitro* translation defect observed for the d15 RNA was rescued by higher levels of RNA synthesis in transfected AC16 cells, resulting in the presence of increased numbers of templates for translation of viral proteins. The increased RNA synthesis directed by the d15 RNA is likely due to the presence of a PCBP2 binding site in stem-loop I of the CVB3 5’ UTR that is missing in the d50 RNA. It has been suggested that these deletions differentially destabilize the secondary structure of stem-loop I RNA but do not completely inhibit PCBP2 binding (29), thereby allowing low levels of viral RNA synthesis. In addition, the altered RNA structures that result from the 5’ terminally deleted genomes may affect the innate immune response to infection (30).

When the biological activities of the different RNA forms (5’ terminally-deleted and full length) were analyzed in mixtures using proportions similar to what was observed in patients suffering from chronic cardiac pathologies [i.e., 19:1 – terminally-deleted RNAs:full-length RNAs], we saw an increase in protein synthesis directed by both forms (Fig. 3C). This underscores why we almost always observed full-length RNA forms in cardiac samples, in addition to the 5’ terminally-deleted RNAs. Although CVB3 genomic RNAs harboring 5’ terminal deletions can replicate and produce higher viral protein levels than forms encoding an inactive 3D polymerase (Fig. 1E), the levels remain extremely low and detection in cell models relies on very sensitive methods. It is possible that full-length forms are needed for efficient replication and to maintain minimal levels of genomic RNA replication. Interestingly, the mechanism was slightly different for RNA synthesis. When mixed in the ratio [19:1-terminally-deleted RNAs:full-length RNAs] as described before, the full-length form had increased RNA synthesis levels, but not the deleted forms. Similar observations were made by immunofluorescence assays targeting an intermediate of RNA replication (dsRNA). Efficient RNA synthesis was highly dependent on an active RNA-dependent RNA polymerase encoded by the full-length RNA form, whereas an inactive polymerase encoded by the deleted form had only a minor impact on dsRNA formation (Fig. 5).

These results highlight the possible interactions of the different forms of enterovirus RNAs (5’ terminally-deleted and full-length genomes); however additional issues need to be addressed. Our cell culture model does not recapitulate any of the immune pressures present during cardiac infections, possibly bypassing the negative selection that may occur when viruses replicate at high level in cardiomyocytes (31). Nonetheless, simulating a re-infection event in a cell culture model (Fig. 6), we were able to detect a significant inhibition in lytic viral particle production, providing a possible explanation for mechanisms used by enteroviruses like CVB3 to persist in cardiac tissue. The exact molecular mechanism(s) remains to be investigated. It has previously been reported that during persistence, enteroviruses exhibit a decreased positive to negative strand RNA ratio, leading to the encapsidation of negative strands as well as positive strands (13). We hypothesize that the deleted forms of enterovirus RNA are able to highjack the cell for virus functions, as evidenced by the cleavage of eIF4G as a marker of host translation shutdown and the formation of dsRNA puncta (observed in immunofluorescence assays) that are normally associated with replication organelle formation and dependent on cellular membrane remodeling. When a re-infection event occurs in these persistently infected cells, there is no lag time during the establishment of viral replication complexes, and large amounts of viral RNA and protein can then be synthesized rapidly. To escape immune surveillance and/or avoid activating a highly inflammatory immune response, we suggest that an enterovirus infection of cardiac tissue is self-limiting, possibly by 5’ terminally deleted genomes causing a defect in lytic viral particle production. Interference with lytic viral production by defective viral genomes has been very well documented in cell culture studies for both RNA and DNA viruses (32, 33), although a connection to human disease pathology has been difficult to ascertain. In addition, complementation of defective viral genomes has been documented during local viral spread for influenza (34), akin to what may occur when there are mixtures of full-length and 5’ terminally deleted CVB3 genomic RNAs in cardiac tissue (10).

The conserved ratio of 5’ terminally deleted genomes to full-length RNAs appears to be important for enterovirus persistence. Non-deleted forms provide proteins to increase the levels of RNA synthesis of deleted forms, improving the overall viral RNA and protein production. However, 5’ terminally deleted genomes inhibit the production of infectious lytic particles. This balance allows for a basal level of viral replication to ensure long-term persistence while preventing a rapid spread of the virus that would result in high levels of immune response and death of the host cell. The exact mechanisms regulating the proportion of full-length to 5’ terminally deleted forms in cardiac tissue remain to be investigated.

To date, no efficient treatments for enterovirus persistent cardiac infections are available. Treatment with interferon β appears to improve outcomes (35) and helps to clear the virus; however, this treatment will not reverse the pathologic damage caused by a long-term, persistent infection. Based on our results and previous studies (20, 21), we suggest that early detection efforts should be coupled with novel therapies targeting the 2A proteinase of enteroviruses. Inhibiting this enzyme would provide at least three beneficial outcomes: (i) shutdown of the processing of the viral polyprotein, thereby inhibiting the infection; (ii) prevention of the shutdown of key cellular process such as cap-dependent translation, allowing for a more efficient innate immune response; and (iii) limiting the pathogenesis linked to 2A proteinase-mediated disruption of dystrophin and other crucial cardiac proteins.

## Acknowledgements

We are grateful to Matthew Marsden for helpful discussions and critical comments on the manuscript. Plasmids harboring the complete CVB3 genome and the associated 5’ terminal deletions of 15, 50 and 100 nucleotides were kindly provided by Laurent Andreoletti (Université Reims Champagne-Ardenne, Reims, France). Research in the authors’ laboratory was supported by U.S. Public Health Service grants AI145003 and AI155962 from the National Institutes of Health (to BLS) and by a postdoctoral fellowship from the George E. Hewitt Foundation for Medical Research (to AB).

